# Computationally mapping olfactory receptors to odor percepts using docking energy scores

**DOI:** 10.64898/2026.06.30.735315

**Authors:** Yuanfang Guan

## Abstract

Mapping the olfactory factors directly to perceived smells has broad implications for neuroscience, chemistry, and medicine. Since the discovery of olfactory genes in 1991, the completion of the olfactory code has been hindered by two obstacles: the unknown combinatorial principles by which ∼400 receptors collectively encode thousands of perceptually distinct odors, and the absence of a complete functional map from the olfactory receptors to conscious perception. We investigated this problem by using the binding energy scores of odorants with the olfactory receptors. We first showed that using only docking scores, we can predict the smell percepts with an accuracy similar to a full set of chemical fingerprints, combining the two resulted in even better performance. This supports there is a direct relationship between olfactory receptors and specific smells. Next, we turned on the olfactory receptors one by one by iterative training and simulation, and produced corresponding specific perception profiles for each olfactory receptor. The generated matrix is sparse, with only 1-2 smell types activated for each olfactory receptors. Despite the limitation of the size of the training data, it suggests the possibility of a low-dimensional combinatorial principle underlying thousands of smells that humans can perceive. We confirmed the prediction by a list of well-known olfactory receptors. We applied the model trained on single chemicals to mixtures, confirming competitive binding was the driving force for smell specificity. This strong performance, surprisingly, is established on the simplest modeling of the binding scores on hundreds of chemicals, and we believe the mapping can become more accurate if more complicated structural modeling techniques and more data are used.

## Introduction

Humans possess approximately 400 functional olfactory receptor genes, each expressed in a discrete subpopulation of olfactory sensory neurons within the nasal epithelium ^1^. These neurons project to spatially invariant glomeruli in the olfactory bulb, forming a topographic map of receptor identity. Yet a fundamental question has remained unanswered for over three decades: how does individual receptor map to specific odor percepts?

Unlike other sensory modalities, olfactory signals reach the cortex without an obligatory thalamic relay, projecting directly from the olfactory bulb to the piriform cortex and higher associative regions, such as the orbitofrontal cortex and limbic structures ^2^. This unique anatomy has hindered the application of general sensory study frameworks to olfaction. For example, electron microscopy-based connectomics maps synaptic architecture ^3^, but cannot resolve functional odor-receptor specificity. Functional MRI captures whole-brain activity but lacks the cellular resolution to resolve individual receptor contributions ^4^. Similar limitations exist for other electrophysiology and calcium imaging techniques. Furthermore, *in vitro* receptor deorphanization often relies on non-physiological, high-concentration ligand screening and often concludes in broad tuning, which can be an inflation from artificial data ^5^. Additionally, most current genetic association studies in humans have primarily focused on olfactory intensity and valence, rather than the genetic basis of specific odor quality perception ^6^. Finally, while many contemporary computational models predict odor percepts from chemical features with impressive accuracy, they remain “black boxes” that fail to map these predictions back to individual receptors. These concurrent failures of experimental and purely descriptive models underscore the need for a mechanistic, receptor-centric computational framework.

The convergence of recent advancements in high-throughput protein docking simulations and the accumulation of large-scale human olfactory perception datasets now provides an unprecedented opportunity. We propose a direct computational mapping of olfactory receptors to odor percepts, bypassing the limitations of traditional physiological screening hurdles. We hypothesized that the high-affinity molecular docking of odorants to olfactory receptors, rather than simply aggregating chemical properties, is the primary determinant of odor specificity. This hypothesis was supported by the strong performance of predicting the smell of odorants based on docking energy scores only, which likely also provides independent information from chemical features. Based on this hypothesis, we computationally mapped the olfactory receptors to specific smells through an iterative machine learning approach. The predictions are strongly supported by published literature, as well as computational control experiments in mixtures.

## Results

### Binding scores predicts olfactory receptor-based odor percepts but not other non-related smell attributes

Before we map the olfactory receptors to individual smell types, we must show that there is a clear predictive relationship between olfactory receptors and odor percepts. We need an intermediate to link the two, because there is no direct training dataset where the two parties co-exist. The chemicals whose odor profile has been described may serve for this purpose. Therefore, for each molecule, instead of fingerprinting-based features, we used a feature vector of binding energy scores against different olfactory receptors. We implemented a specialized computational architecture that allowed us to generate the score of each ligand against each olfactory receptor.

We used the DREAM Olfactory Mixture Dataset downloaded from Sage Bionetworks Synapse ^7^, where each molecule has 54 smell types (including intensity and pleasantness) labeled by quantified numbers. This dataset was originally developed for another purpose: benchmarking machine learning algorithms to predict smells based on chemical structures, but we can leverage it for our purpose. Several alternative datasets are expected to yield comparable results. We chose this one because of some of its additional information, *e*.*g*., similar/repetitive descriptors, intensity (which should not be predicted by docking), existence of non-olfactory receptor-associated smell types and chemical mixtures, which can help us develop many sanity checks throughout the process. For the purpose of accurate mapping, other larger datasets will be more preferable.

For the docking scores, we implemented a decoupled computational architecture that pre-calculated AlphaFold3 ^8^ structural profiles and optimized binding energy (ΔG) grids using Vina ^9^, allowing us to perform large-scale virtual screening of ligand-receptor interactions. Clearly our approach suffers from the inaccuracy from rigid modeling. There are other more accurate approaches such as Alphafold3 or molecular dynamics refinement to further improve performance, but our approach helped to overcome computational bottlenecks and allowed us to iterate the experiments quickly. We reasoned that a genuine receptor-percept relationship should be detectable even under conservative docking assumptions.

We carried out five-fold cross-validation to examine whether the feature vector of binding scores can predict the smell types. For each smell type, we trained a lightGBM model. Using LightGBM might involve a potential pitfall: while lightGBM does not explicitly force sparsity like L1 regularization, it tends to select the important features. Then how can we be sure that the observed sparse mapping of an olfactory receptor to one or two specific smells is true observation or an artifact of LightGBM? Note that the model was trained for each smell type, instead of for each receptor. Therefore, if an olfactory receptor is very important for multiple smell types, we will still be able to pick it out. Indeed, this was observed for closely related terms as will be shown soon. Alternative tree-based algorithms will serve the same purpose well, which is beyond the scope of this study.

We found that other than intensity, sharp, ammonia, or alcoholic, the model can successfully predict the smell type simply based on the binding profiles (**Figure 1**), indicating that there is a direct relationship between the olfactory receptor binding and specific smell type. Intensity is not supposed to be predicted well, since no concentration data was input into the model, while the chemical intensities were diverse. Sharp and ammonia are not perceived by olfactory receptors. They are mediated by the trigeminal nerves^10,11^ . The model’s inability to capture these non-olfactory or concentration-dependent attributes serves as a sanity check.

**Figure 1.**
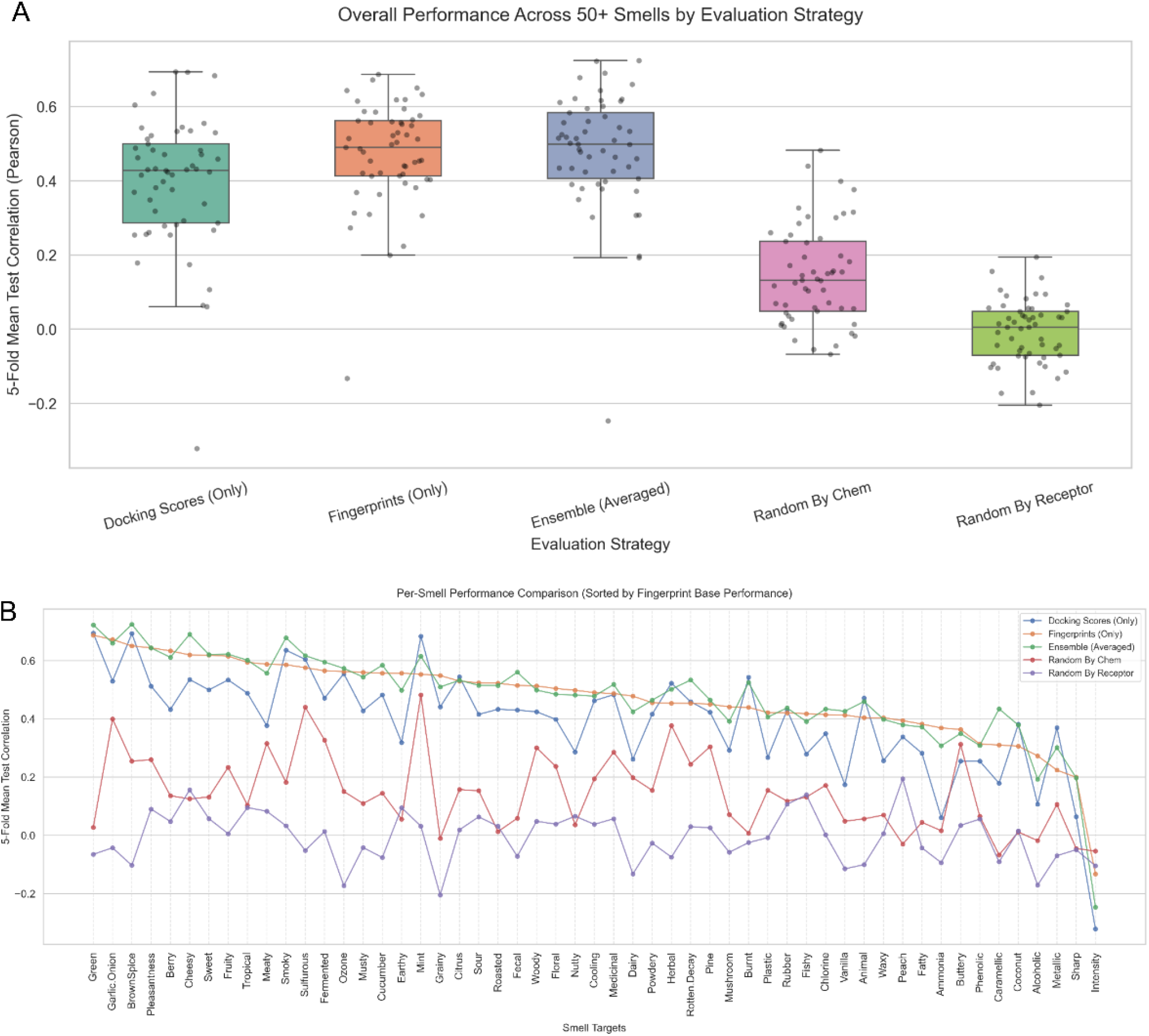
Overall performance comparison across different methods. A. Box plot of Pearson correlation coefficients across 52 smell descriptors, pleasantness, and intensity. *Docking score*: docking energy scores generated by AlphaFold3-Vina used as features. *Fingerprinting*: ensemble of four chemical fingerprinting feature sets. *Ensemble*: average of predictions from the docking score and fingerprinting models. *Random by chem*: docking features randomly permuted for each chemical. *Random by receptor*: docking features randomly permuted for each receptor in the training set. B. Per-descriptor performance comparison across the five models.

### Control experiments with shuffled data and fingerprinting models

Our next question is how much of the information comes from the fact that a particular chemical has broad interactions (as suggested in *in vitro* screening experiments ^12^), versus how much it comes from the specificity of olfactory receptors. For each chemical, we randomized the olfactory receptor order in training and testing. The resulting median correlation is 0.1317 compared to 0.4279 from the original model. This result indicates that the receptor-specific information is the main driving factor for predicting the odor percepts. As a negative control, we randomized the chemicals for each olfactory receptor in the training set, the resulting median correlation is 0.0058, essentially random.

To get a sense of the meaning of this performance, we carried out the following comparison experiments. First, we used four chemical feature sets that vary by number of features and types as training features: molecular descriptors (5 basic structural and physicochemical properties), Molecular ACCess System keys (MACCS, a structural fingerprint with 166 bits), Morgan (circular fingerprints, with 2048 bits) and RDKit-generated fingerprinting (linear substructural paths, with about 2000 features). Surprisingly, docking scores alone performed similarly to all molecular features combined together using the exact training parameters (**Figure 1A**), and outperformed Morgan and Descriptor molecular feature sets (**Figure S1-2**). We ensembled the predictions of the two models together, which slightly improved the overall result. The performance of the fingerprinting model has a correlation of 0.7325 with the docking model, indicating that the list of odorants existing in the training set is the main limiting factor to model performance. The only exceptions are ammonia and sharp, where the docking model performed substantially worse than the fingerprinting model. This makes sense because ammonia is not a smell type sensed by the olfactory receptors, confirming the absence of implicit chemical information leakage in the docking features.

### Theoretical activation analysis maps olfactory receptors to a percept profile

To establish a biological reference for the predictive model, a resting background profile was established using precalculated median binding energies for each olfactory receptor across all testing chemicals. On the other hand, the maximal “Turn On” or activation threshold for each specific receptor was defined as the absolute minimum energy value observed across the training matrix. The distribution of these maximal “Turn On” values ranges from -5 to -12 kcal/mol, with the majority centered at -9, indicating moderate to strong bindings which is reasonable for our simulation, compared to the overall binding score distribution which is centered at -4.7 kcal/mol (**Figure 2A-B**). On average, each olfactory receptor has 2.28 molecules with a strong hit to it.

**Figure 2.**
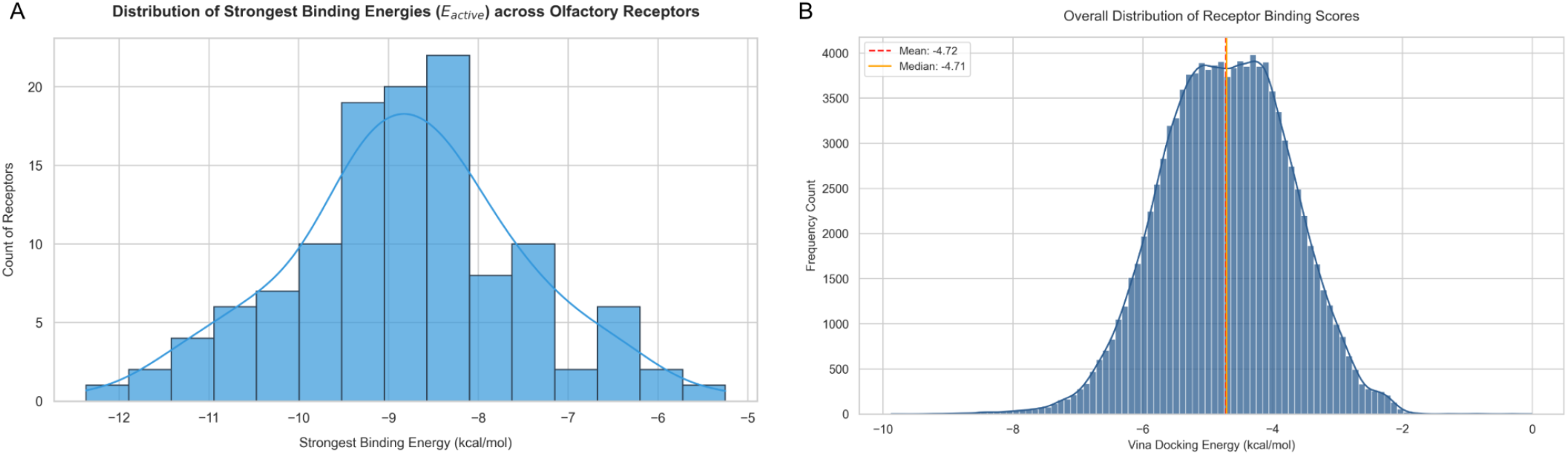
Distribution of binding energy. **A**. For each olfactory receptor, we identified the lowest score of binding energy across all chemicals and plotted this distribution. **B**. Distribution of all binding scores over all chemicals and over all olfactory receptors.

To evaluate the direct contributions of individual olfactory receptors to specific percept, a one-at-a-time *in silico* activation simulation was conducted using pre-trained LightGBM models combining all molecules. For each target smell profile, an initial baseline prediction score was generated by passing the resting median background profile through the corresponding LightGBM regressor. Receptors were systematically activated in isolation to quantify their functional contribution. For each individual receptor *i*, the feature vector was duplicated, and its baseline value was substituted with its corresponding activation value (the lowest binding score in the training data). The modified feature vector was then passed through the LightGBM models to generate activated prediction scores. The absolute impact of receptor *i* on a specific odor descriptor was calculated *via* a delta-subtraction framework.

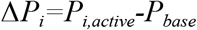

A positive *ΔP*_*i*_ indicated a promoting effect of the receptor on the perceived smell type, whereas a negative *ΔP*_*i*_ signaled an inhibitory effect.

The model for each smell type was trained independently, yet the signal is sparse for all olfactory receptors. The only exceptions are smell types that are intrinsically related to each other, *e*.*g*., mint and herbal, and another example is fruit, berry, peach (**Figure 3**). They are often predicted together. This observation is true for the entire olfactory receptor family we examined. Whether this sparsity reflects a true property of olfactory coding or an artifact of the limited training set size remains an open question, which is addressable by applying the same framework to larger chemical datasets.

**Figure 3.**
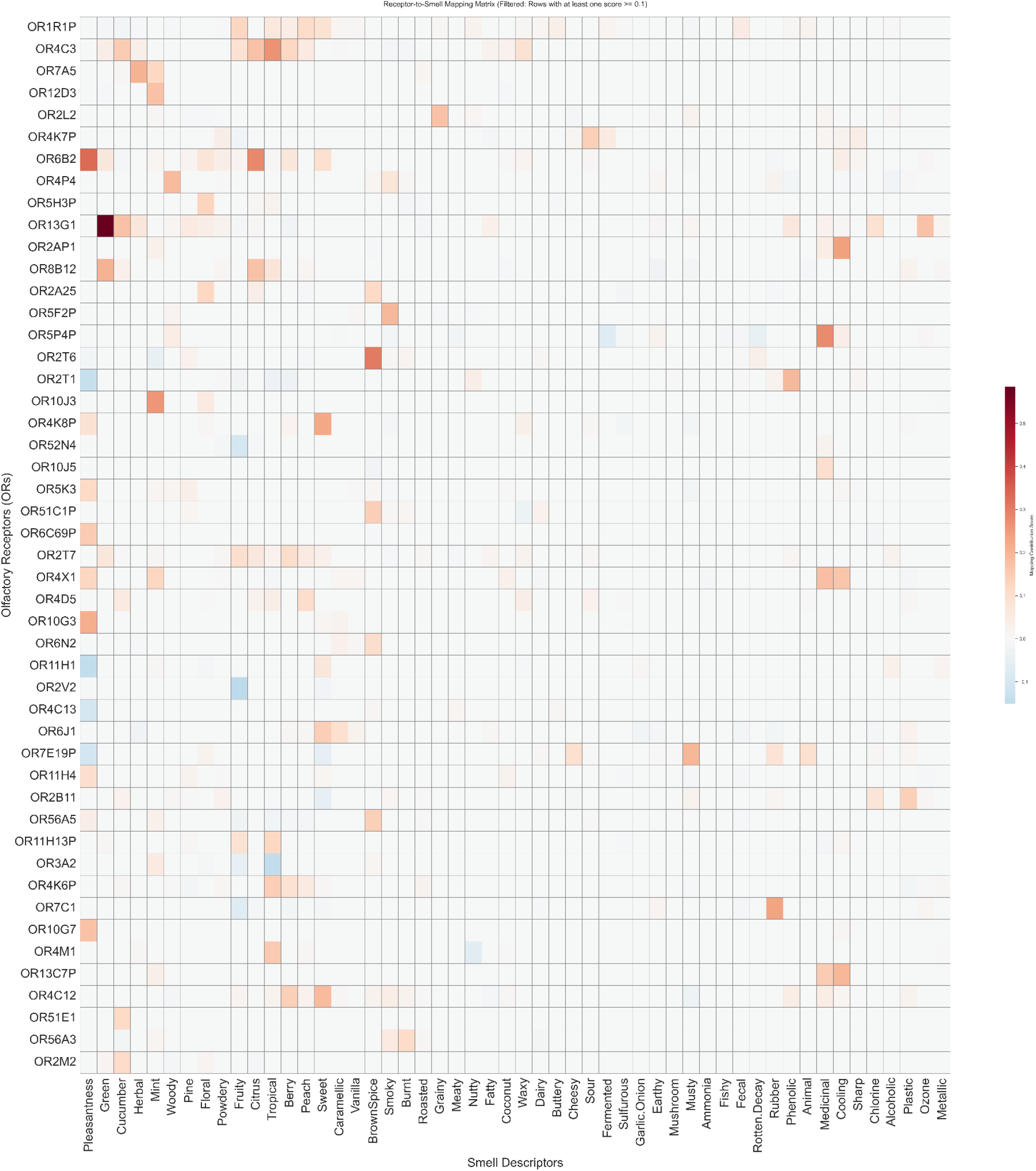
Baseline round activation analysis maps olfactory receptors to percept profiles. The heat map displays the calculated activation impact of individual ORs across diverse semantic smell types. Each row represents a specific OR, and each column represents a targeted odor percept; color intensity denotes the magnitude of the shift in predictive probability relative to the baseline.

### Iterative multi-round computational knockout to predict odor percepts of more olfactory receptors

Because olfactory receptors frequently possess overlapping, highly co-linear pocket geometries, tree-splitting algorithms may isolate a single dominant feature and ignore its biological twins. To circumvent this “all-or-nothing” selection, we implemented an iterative feature knockout pipeline. Let R_0_ be the initial pool of all available olfactory receptors. In any given iteration *t*, the active receptor feature pool R_t_ was defined by cumulatively removing all dominant receptors (those with an activation impact ΔP >0.1) identified in previous rounds. This allowed us to predict 30-50 olfactory receptors at a time. For each iteration, we first carried out five-fold cross-validation with the leftover features to monitor if the model is still performing reasonably. We found that at least in the first 10 iterations (**Figure 4**), we obtained reasonable performance. This implies that either some of the olfactory receptors are redundant or there is a limitation in olfactory vocabulary in describing the smells. Then, a final model was built on all chemicals to identify receptor-associated odor type and for removal in the next iteration. The first 10 iterations assigned 391 out of 516 receptors to specific smell types.

**Figure 4.**
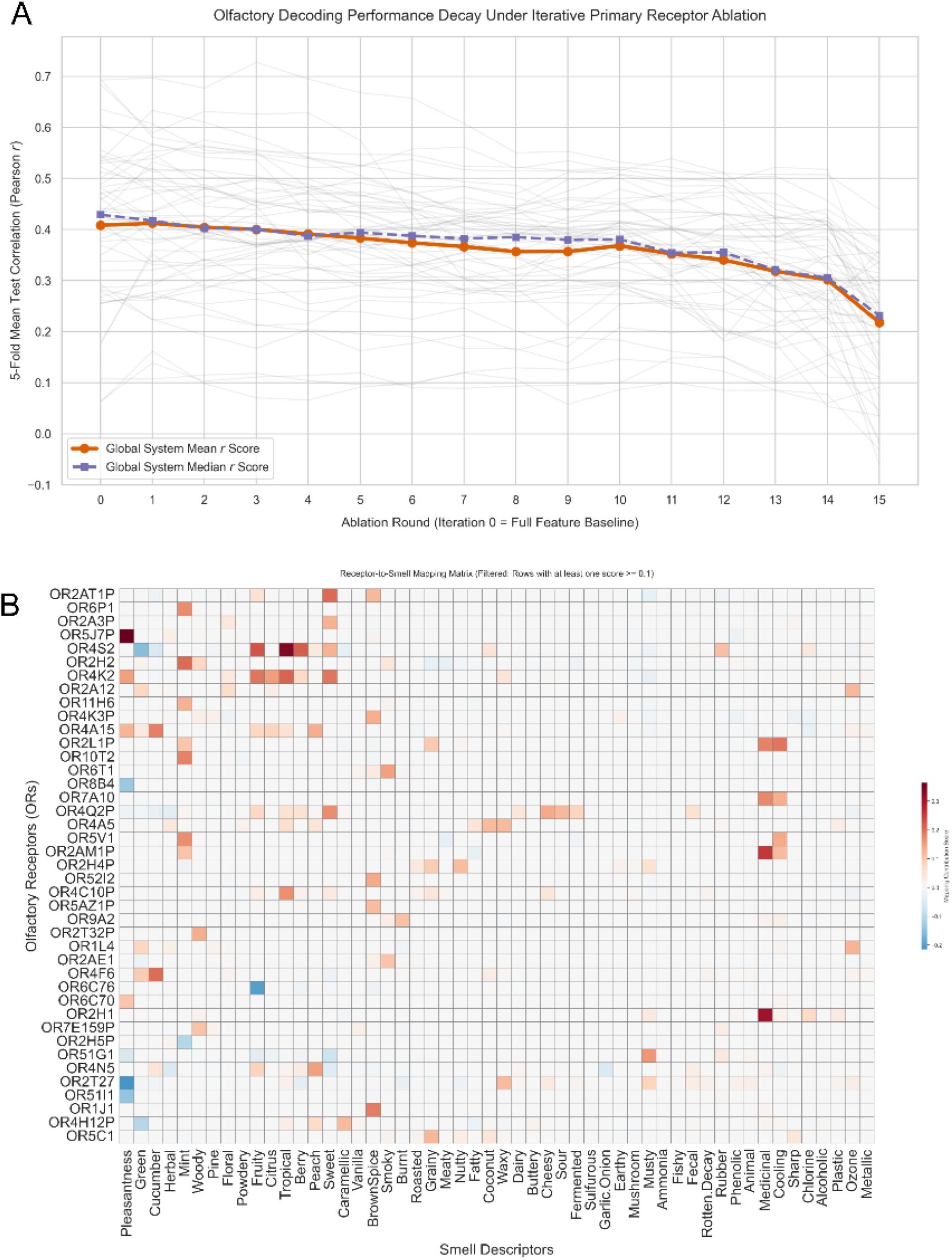
Iterative strategy to predict the corresponding smells of additional olfactory receptors. **A**. Performance changes across iterations. Later round predictions are less reliable. **B**. Round 1 prediction matrix.

We identified six receptors in literature where there is existing evidence of what smell type they are associated with. We saw strong consistency between the predicted olfactory type and the ones known in literature (**Table 1**). For example, the model predicts that OR10G3 corresponds to pleasantness and sweetness. OR10G3 is a known receptor to vanillin ^13^, which is the gold standard for high pleasantness, sweet, and caramellic smells. OR2M2 is selectively activated by the structurally distinct ligands 1-p-menthane-8-thiol and nootkatone, both of which are associated with grapefruit aroma; our model instead predicted OR2M2 to be associated with cucumber/green descriptors, which is a partial but biologically plausible match, given the shared fresh, green-toned quality of these percepts ^14^. There are also cases of suppression, the implication of which will be discussed later.

**Table 1.** Example predictions compared to physiology study in literature. (-) means suppression effect.

| Receptor | Primary Literature Profile | Top Predicted Descriptor | Activation Round | Interpretation |
| --- | --- | --- | --- | --- |
| OR10G3 | Vanillin (Vanilla) | Pleasantness | Baseline (0) | Capturing highly pleasant vanilla/sweet motifs. |
| OR2M2 | Aliphatic Aldehydes | Cucumber | Baseline (0) | Capturing green, cucumber food volatiles. |
| OR7D4 | Androstenone (Steroid) | Green (-) / Brown Spice (+) | Round 2 | Structural modeling of heavy warm woody/spice steroid tones. |
| OR2J3 | Leaf Alcohol (Cut Grass) | Mint | Round 2 | Transition of fresh green aliphatics into the mint boundary. |
| OR1A1 | Terpenes & Cyclic Alcohols | Medicinal (+)/ Pleasantness (-) | Round 5 | Capturing the sharp, industrial, un-pleasant terpene threshold. |
| OR51E2 | Short-chain Fatty Acids | Musty / Waxy / Earthy | Round 6 | Broader mapping of organic fermentation and waxy lipid states. |

### Predicting odor profile of mixtures by using the strongest binding value across all components

Although our model was trained on single chemical compounds, we can use it to test the following hypothesis: if specific binding affinity is the primary determinant of olfactory perception, selecting the minimum energy score across all constituent chemicals in a mixture should yield better model performance compared to utilizing the mean or maximum values. Of note, this approach is not intended to optimize prediction performance, but rather to validate our overall hypothesis underlying the assignment of olfactory receptors to smells.

We observed that when taking the lowest energy score, the median correlation is 0.190, while using the maximal score had a median correlation of 0.117 (min versus max Wilcoxon test, *p* = 0.00056), and using the mean score across all chemicals had a correlation of 0.134 (min versus mean Wilcoxon test, *p* = 0.0047, **Figure 5**). The superior performance when using strongest predicted docking affinity suggests that olfactory perception is governed by a ‘winner-take-all’ mechanism at the receptor level. In this framework, the constituent ligand with the highest affinity for a specific olfactory receptor acts as the primary agonist, effectively driving the signal cascade and defining the perceived odor. These results support the hypothesis that specific, high-affinity molecular docking is the key determinant of receptor-mediated odor discrimination, rather than an aggregate mixture effect.

**Figure 5.**
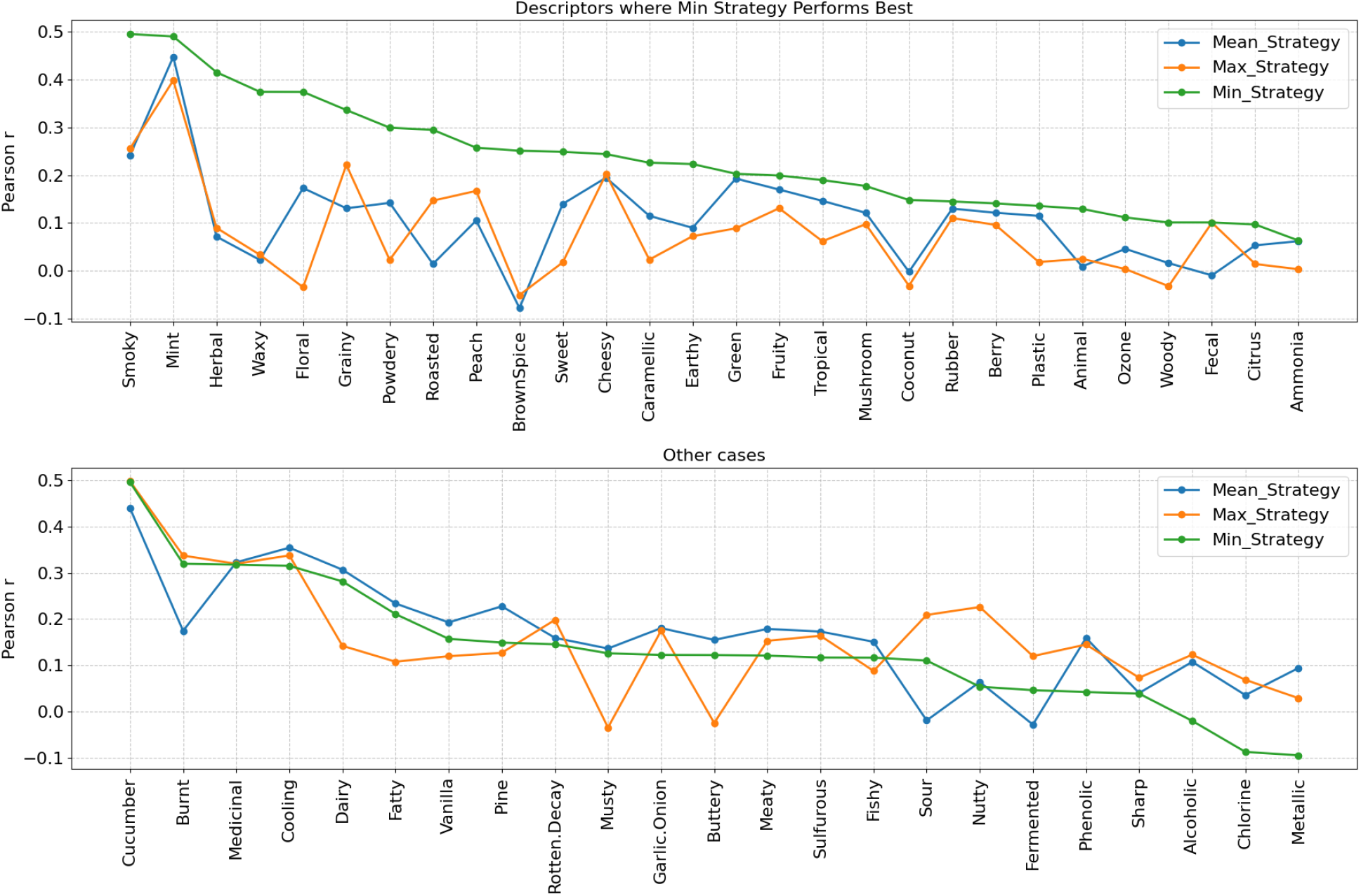
Combinatorial binding profile predicts the odor profile of mixtures when using the strongest binding value (lowest score). For many smells the min scores significantly outperformed the other methods in cross-validation.

## Discussion

This study demonstrates that molecular docking scores between odorants and olfactory receptors carry sufficient information to predict human odor percepts, establishing a direct computational link between receptor-level binding and perceptual outcomes. It also left open questions due to the limitation of the dataset we used in this study. We will discuss them one by one.

It is confirmed that simple binding scores can predict the smell percepts, even when the binding scores were inferred roughly using computationally simplified yet efficient methods, and even when the base learner was simple and not optimized. We implemented many sanity checks throughout the experiments to confirm this point. We showed some empirical, computational and logical evidence that the one-at-a-time simulation might reveal how the olfactory receptors are related to each smell type. It is observed that the most commonly associated smell we found was fruit/sweet. However, in this case, we do not have any clue if it were an artifact due to good performance (either easier to predict or more training examples), or evolutional benefit, or both.

We observed that the encoding learned in this study is sparse. Certainly with a sparse encoding, combinatory effects can produce many more smell types that humans can differentiate. However, it is also possible that this is an artifact due to the small training set, from which we do not have enough strong hits for each olfactory receptor. Training with a large dataset is expected to answer this question.

We do not fully understand the few suppression cases that either come from model noises or underlying mechanisms, where the model predicts a negative value. For example, the only strong prediction we made for the orphan receptor OR6C76 is suppressive behavior on Fruity. We evaluated this prediction by isolating the most potent stimuli (top 10%) of fruity and earthy respectively. We observed that “Earthy” odorants exhibit a significantly higher binding affinity to OR6C76 compared to “Fruity” odorants (*p < 0*.*002*). This thermodynamic preference suggests that OR6C76 functions as a competitive filter: potent Earthy molecules occupy the receptor binding pocket with greater stability, effectively out-competing Fruity ligands. This mechanism suggests a structural basis for the behavior of mixture suppression, where earthy signatures are known to dampen the perception of delicate fruity notes. The observation suggests that OR6C76 is not merely as an orphan receptor, but as a “suppressive module” within the olfactory epithelium, maybe serving to refine odor discrimination by filtering out high-valence fruity signals in the presence of dominant environmental earthy backgrounds. This observation is consistent over all of the remaining suppressive behavior we identified of fruity in the Baseline round and Round 1: OR52N4 (*p* =0.0006), OR2V2 (*p =* 0.0011), OR7C1 (*p* = 0.0014). Systematic analysis over all suppression cases with a large collection of chemicals can potentially confirm whether the competitive binding mechanism is an artifact or true phenomenon.

There are many areas of future improvement for this study. We used a tiny training set. If more training data is involved, we expect better performance and clarification of some of the above questions. There are several smell types, vanilla, sulfurous, mushroom, fishy where none of the olfactory receptors showed a strong association. This is likely to be an artifact of lack of training data, as they performed poorly with the fingerprinting model too. The only exception is ammonia (or sharp), where the fingerprinting model performed well but not the docking model. This is likely due to ammonia not “smelled” with standard olfactory receptors, but with trigeminal nerve system. Similarly, more accurate docking analysis could further improve the results, for which there is a considerable room for improvement. Several methodological limitations of the computational screening pipeline warrant acknowledgment. First, affinity grid placement was anchored to the geometric centroid of the full receptor structure rather than an experimentally validated or cavity-detected binding site. While this approach is acceptable for large-scale automated screening of structurally diverse OR models, it introduces positional uncertainty in cases where loop length, terminal tail geometry, or AF3 prediction artifacts displace the whole-protein barycenter from the true orthosteric pocket. Future work could address this by anchoring grids to conserved functional residues or by incorporating an upstream cavity detection step using tools such as fpocket ^15^ or DoGSiteScorer ^16^. Second, only the single top-ranked docking pose was retained per receptor-odorant pair. This approach maximizes throughput, simplifies the downstream machine learning model but sacrifices information about binding mode degeneracy. An alternative approach is to test a clustered ensemble of low-energy poses. Third, hit prioritization relied exclusively on AutoDock Vina’s empirical ΔG scoring function. While sufficient for this explorative study, this scoring function has known limitations for GPCR targets, particularly in polar or charged binding environments. Rescoring top candidates with a complementary approach, such as the Gnina CNN-based scoring function or an MM-GBSA free energy approximation^17,18^ would improve confidence in the final hit list. These limitations are consistent with the scope of a high-throughput virtual screening study. Addressing them in future work would likely improve docking accuracy and hit confidence; however, they fall outside the primary objective of this study, which is to develop an initial machine learning framework for mapping olfactory receptors to individual smell categories.

## Methods

### Decoupled computing architecture for molecular docking

Processing each receptor-ligand pair end-to-end through AlphaFold3^19^ alone is computationally prohibitive at scale. We therefore decoupled the resource-intensive evolutionary MSA construction step from the downstream docking screen. The structural search profile for each receptor was computed only once and cached, rather than recomputed for every receptor-odorant pair. This reduced the combinatorial compute cost from O(M × N) (M receptors × N odorants) to a linear cost in M, making large-scale screening computationally feasible.

Multiple sequence alignment (MSA) profiles encoding evolutionary conservation were precomputed for each target olfactory receptor (OR) using AlphaFold 3 (AF3). Each receptor sequence was submitted as a protein-only JSON input file to the AF3 pipeline, which queried UniRef90 ^20^, MGnify ^20,21^, and environmental metagenomic databases via HMMER ^22^ and HH-suite ^23^. The resulting MSA profiles and associated feature representations were cached to build a reusable structural database for downstream docking.

Three-dimensional receptor coordinates were obtained from AF3-generated mmCIF structure files. To exclude low-confidence or poorly resolved models, each receptor structure was screened by computing the proportion of unidentified or poorly resolved residues relative to total sequence length; receptors exceeding a 5% threshold were excluded from further analysis.

Receptor models passing this quality filter were converted from mmCIF to standard PDB format using Biopython’s PDBIO module. Structures were then processed using Meeko (v0.7.x) to remove non-polar hydrogens, assign Gasteiger partial charges, and define atom types, producing rigid receptor structures in AutoDock PDBQT format for subsequent docking ^9^.

### Structural preparation and dynamic grid optimization of olfactory receptors

High-throughput virtual screening (HTVS) was conducted against the full panel of olfactory receptor (OR) structural models. Because AlphaFold3-predicted structures lack a fixed reference orientation, the docking search box was anchored individually for each receptor at its own structural centroid, rather than using a fixed global coordinate frame. For each receptor, the centroid *C*_*dynamic*_ was computed as the arithmetic mean of the Cartesian coordinates of all atoms (ATOM and HETATM records) in the corresponding PDBQT file:

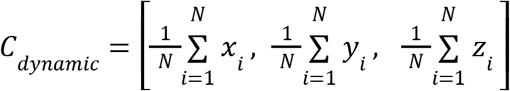

where N is the total number of atoms in the receptor’s PDBQT structure. The affinity grid was centered on *C*_*dynamic*_ using a cubic search box of 25.0 × 25.0 × 25.0 Å, sized to encompass the structural core of the 7-transmembrane barrel and capture odorant interactions within this region. To evaluate the sensitivity of this choice, we additionally tested 20 × 20 × 20 Å and 30 × 30 × 30 Å boxes; docking performance was comparable across all three sizes, with the 25 Å box providing a robust balance between spatial coverage and computational cost (**Figure S3**).

### Predicting sensory profiles using molecular docking scores

To predict human olfactory sensory profiles from molecular docking scores, we implemented a machine learning workflow utilizing LightGBM regressors. The feature matrix *X* was constructed by aggregating molecular docking scores across multiple olfactory receptors (ORs). For each molecule, docking energies against a panel of individual receptors (extracted from Autodock Vina simulations) were concatenated. The background baseline calculated from the median values of each receptor’s scores across all chemicals was saved for downstream normalization during silico activation simulation. Human sensory perception intensities across distinct descriptors were used as the target labels *y*.

A separate predictive model was trained for each individual smell target. To evaluate model generalizability, we employed a 5-Fold Cross-Validation (CV) scheme with a fixed random seed to ensure deterministic sample partitioning. For each fold, a LightGBM Regressor (100 estimators, learning rate = 0.05, and a shallow maximum depth of 2 to mitigate overfitting) was fitted on the training split (80% of the data). Hyperparameters were not extensively tuned; default values informed by prior experience were used, as hyperparameter optimization was not the focus of this study. The performance of the models across each fold was evaluated using the Pearson correlation coefficient between the true human sensory ratings and the model’s out-of-fold predictions.

