## Additional Figures for "Computationally mapping olfactory receptors to odor percepts using docking energy scores"

**Figure S1**. Similar performance of docking scores compared to individual molecular fingerprinting feature set.

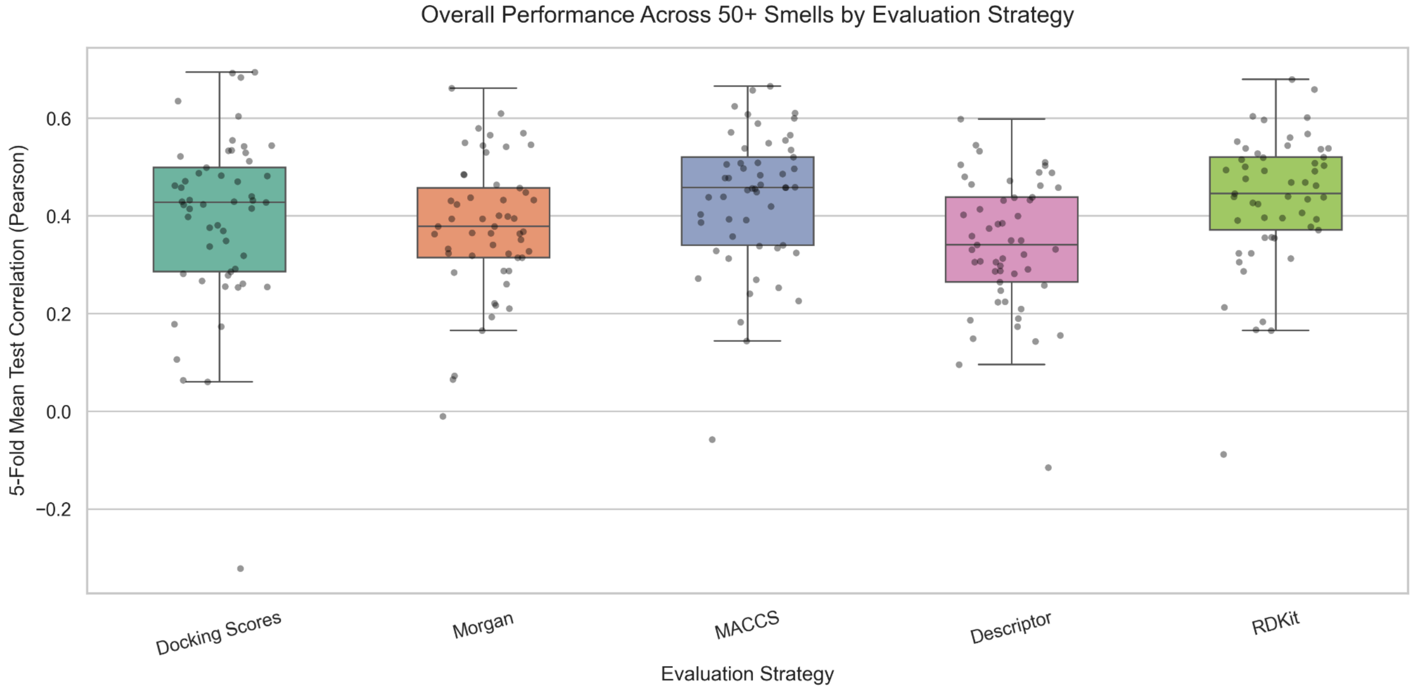


**Figure S2.** Comparison of the performance of docking scores and individual molecular fingerprinting feature set across 52 smell types, pleasantness and intensity.

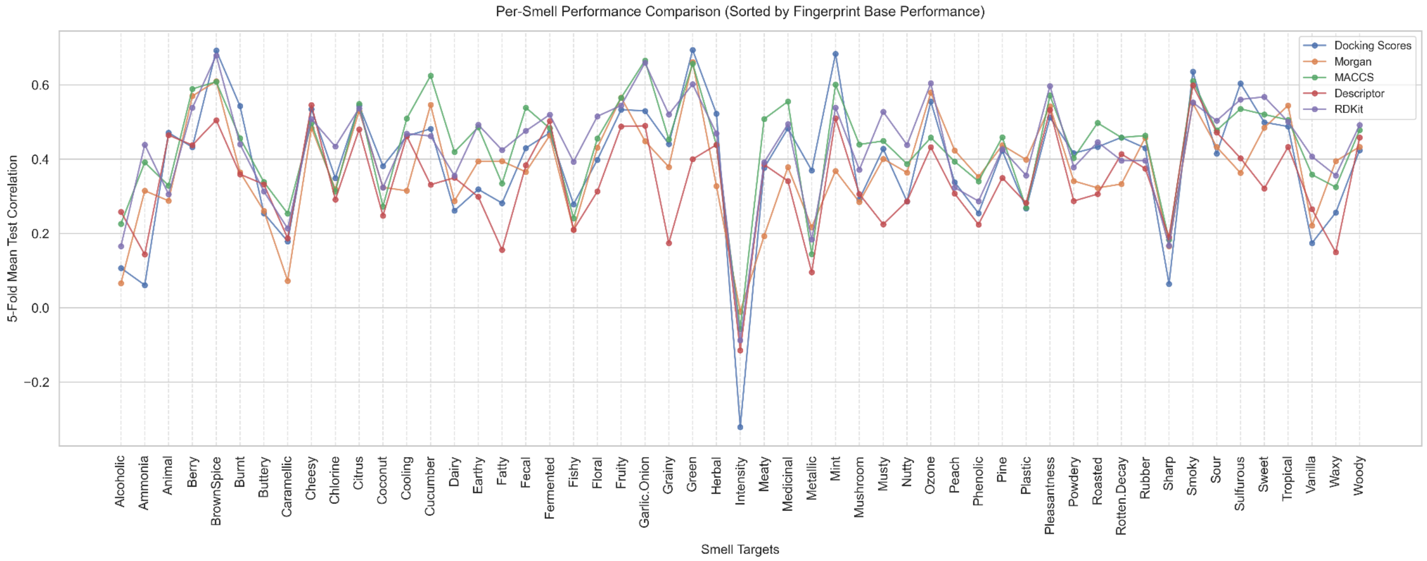

**Figure S3.** Performance is similar across all three cubic search sizes.

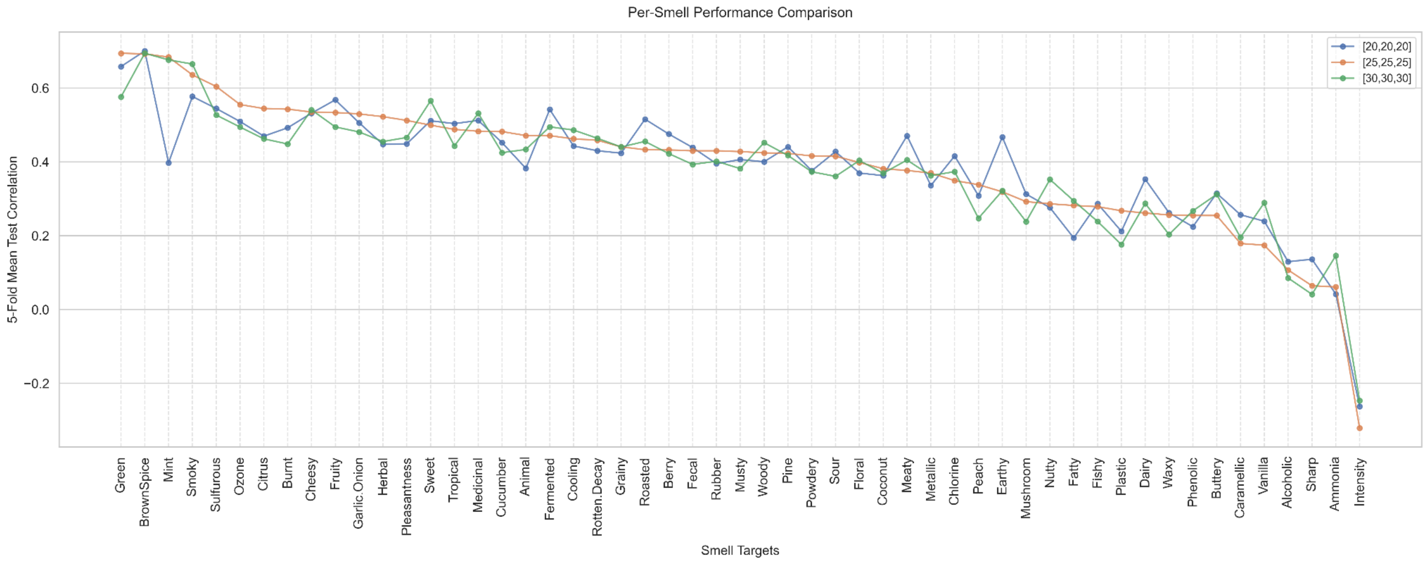
